# Sequence biases in CLIP experimental data are incorporated in protein RNA-binding models

**DOI:** 10.1101/075259

**Authors:** Yaron Orenstein, Raghavendra Hosur, Sean Simmons, Jadwiga Bienkoswka, Bonnie Berger

## Abstract

We report a newly-identified bias in CLIP data that results from cleaving enzyme specificity. This bias is inadvertently incorporated into standard peak calling methods [1], which identify the most likely locations where proteins bind RNA. We further show how, in downstream analysis, this bias is incorporated into models inferred by the state-of-the-art GraphProt method to predict protein RNA-binding. We call for both experimental controls to measure enzyme specificities and algorithms to identify unbiased CLIP binding sites.

---

The peak-calling process in CLIP experiments derives peaks from raw sequences [1]. Bound RNAs are cleaved by RNase T1, which cuts at accessible G’s; as a result, a majority of called peaks terminate at a G (Figure 1A). To demonstrate this, we analyzed CLIP data used in the GraphProt study [2] (see Supplementary Methods) and found the presence of the ‘terminating G’ effect. For each sequence in the peak and control sequences, we calculated the frequency of G’s at the last position. We found that there is a much higher frequency of G’s at the last position of the peak sequences as opposed to in the control sequences (Figure 1B). In the most extreme case, more than 90% of the last nucleotides in the peak sequences are G’s, as compared to 25% in the control sequences. In contrast, when we analyzed the raw sequencing data and its nucleotide frequencies, we did not observe a ‘terminating G’ effect (e.g., Figure 1C); suggesting that this bias is introduced in the peak-calling. In addition to the ‘terminating G’ bias, we also observed a ‘G-depletion’ bias in the peaks, in concordance with Kishore *et al.* [3] (Supplementary Figure S1A). Complete results are in Supplementary Table S1.

**Figure 1.**
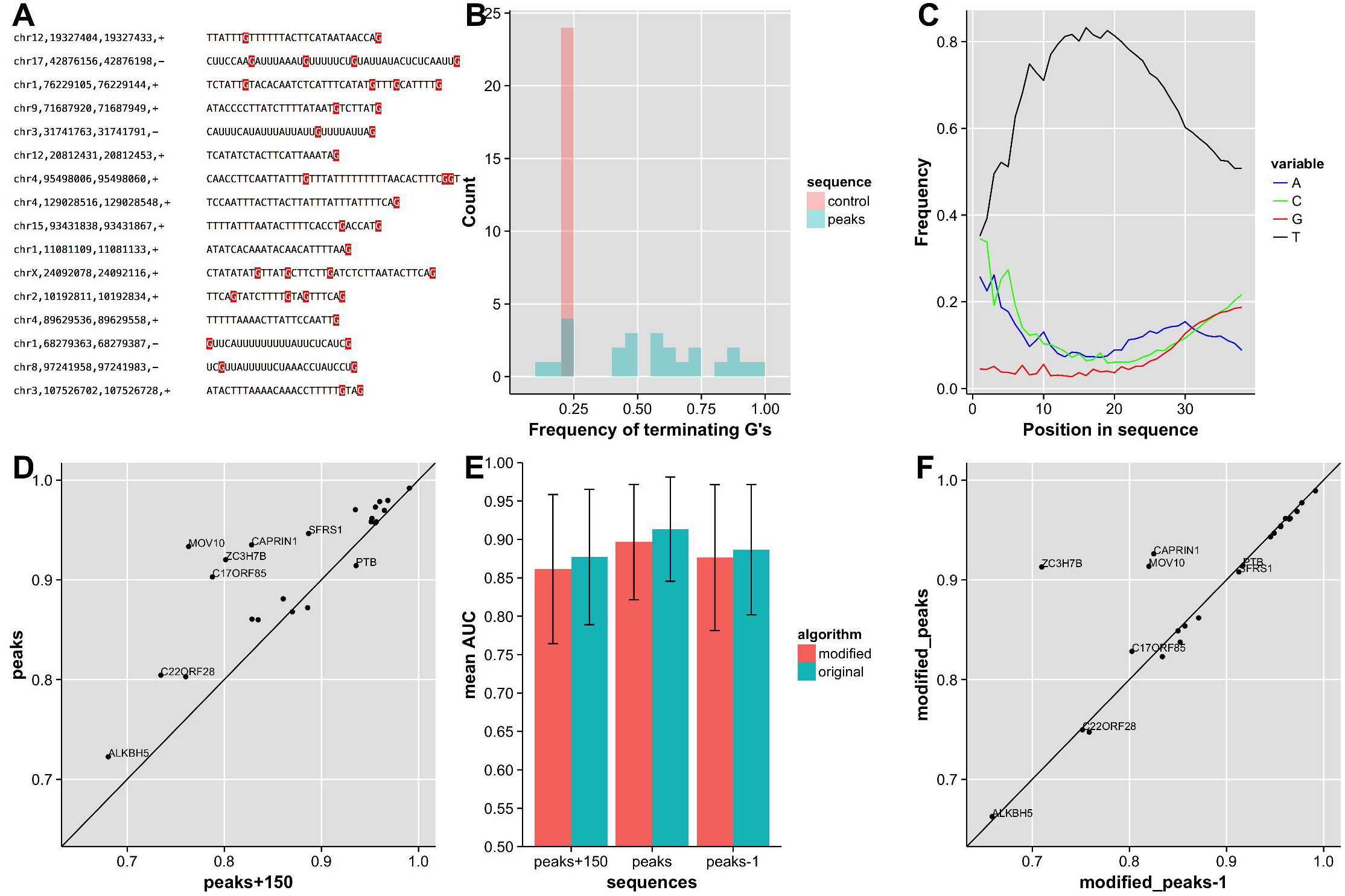
Sequence biases in CLIP experiments and their effect on binding models learned in downstream analysis. A) Most binding site peaks from Elavl1 CLIP-seq experiment terminate at a G (highlighted in red). B) Frequency of terminating G’s is much higher in peaks than in control sequences. The histogram is over the frequency of sequences ending with G’s. C) Most raw sequences do not terminate with a G, indicating the bias is introduced in the peak-calling. The nucleotide frequencies are per position of raw sequences from Elavl1 CLIP-seq experiment. D) GraphProt performance on CLIP experiments without flanks is much better due to terminating G’s being more visible (see Table 1). E) Without the features encoding terminating nucleotides, GraphProt performs worse, but not a worse as not having the terminating nucleotide in the data (*peaks-1*). The original and modified GraphProt performance on different flanks lengths is averaged over 24 CLIP experiments. F) For some experiments, GraphProt modified algorithm benefits from the terminating G even without the terminating features. For other experiments, the performance decreases to the level of not having the last nucleotide in the data (*peak-1*).

To investigate whether this bias is incorporated in computational models, we trained the state-of-the-art method for protein RNA-binding prediction, GraphProt [2], on their original data with three different flanks (denoted by *peaks ± length*): peaks flanked by 150bps– *peaks+150* (as used in the original GraphProt study); peaks alone– *peaks*; and peaks without the last nucleotide– *peaks-1* (see Supplementary Methods for details and justification). We tested the models through 10-fold cross-validation, as in Maticzka *et al*. [2]. We found that models which incorporated the terminating G’s improved their prediction accuracy, an improvement that was even more pronounced without the flanks (Figure 1D). The average AUC was 0.913 for *peaks*, as compared to 0.877 for *peaks+150* (with a p-value = 0.00013, determined by the Wilcoxon rank-sum test, comparing results on 24 CLIP experiments). When we removed the last nucleotide from the peaks, the accuracy dropped significantly (Figure S1B); the average AUC for *peaks-1* was 0.887, as opposed to that of 0.913 achieved with *peaks* (p-value = 0.00087, using the same p-value test here and henceforth). When we investigated the underlying causes of this discrepancy, we observed that many of the highly weighted GraphProt features were encoding terminating G’s (Table 1). Remarkably, when we based our predictions solely on whether the sequence had a terminating G, we were able to achieve higher AUCs than that achieved by GraphProt for 5 (out of 24) experiments (Supplementary Figure S1C). Complete results are in Supplementary Table S2.

**Table 1.**
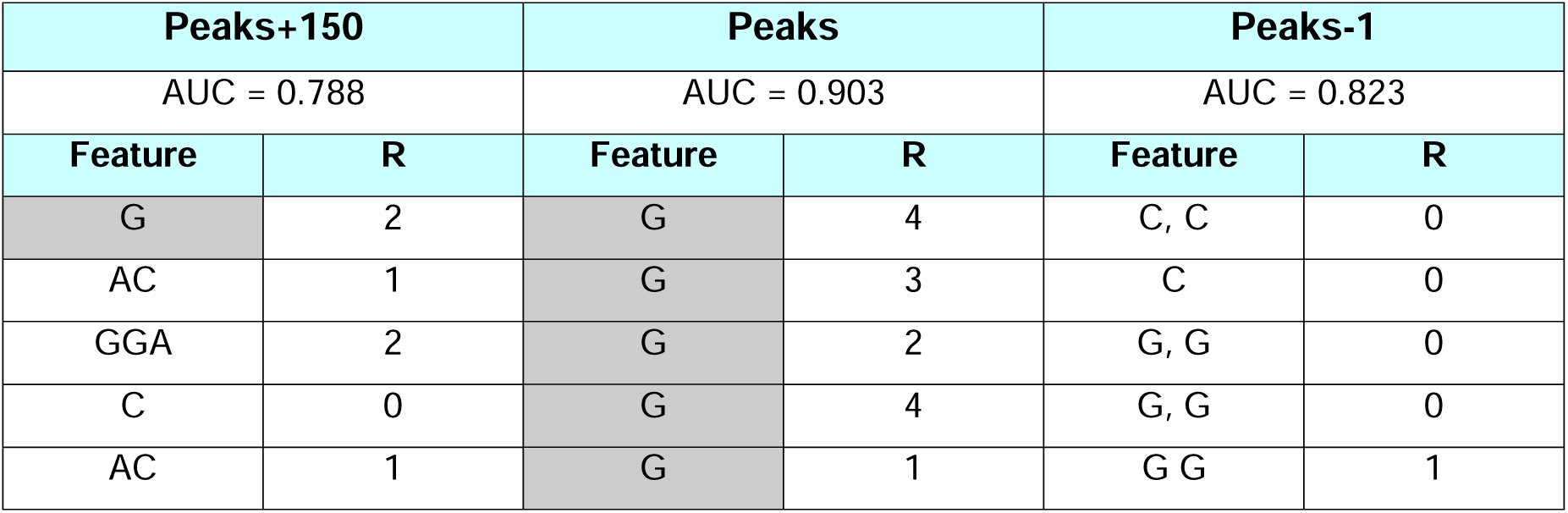
AUC results and top features learned by GraphProt for protein C17ORF85. For three different flank length: peaks and flanks (*peaks+150*), peaks only (peaks) and peaks only without the last nucleotide (*peaks-1*), we used GraphProt to train a model (see Supplementary Methods). We report the accuracy of the models (gauged in 10-fold cross validation) and the top 5 features. For each feature, we report the nucleotide composition and the R parameter by which it was generated. R is the neighborhood radius reachable from the nucleotide and thus G features with R>0 (grey background) encode a terminating G (for detailed feature encoding, see Maticzka *et al.* [2]).

To resolve the effect of this bias on inferred binding models, we modified the GraphProt algorithm to exclude features that encode terminating nucleotides (Supplementary Methods). We ran the modified version on the different flank lengths, as before, resulting in six combinations of algorithm version and flank length. As expected, the performance of the modified version decreased significantly as compared to the original algorithm: the average AUCs for *peaks+150*, *peaks*, and *peaks-1* were 0.870, 0.891, and 0.877, as compared to 0.877, 0.913, and 0.886 for the original feature set, respectively (p-value = 2.3⋅10^−7^) (Figure 1E). Surprisingly, the performance did not decrease significantly when the last nucleotide was removed: the average AUC for the modified version was 0.891 on *peaks* versus 0.877 on *peaks-1* (p-value = 0.002) (Figure 1F; Figure S1D for a comparison of *peaks+150* to *peaks*), implying that the modified version still captures the terminating G bias, albeit with less efficiency. To summarize, GraphProt’s performance decreased without having the terminating features, yet the remaining features still captured some of this bias.

## Conclusions

Here we reported a newly identified bias in the CLIP analysis pipeline. The source of the technological bias comes from enzyme specificity [3], although it is truly introduced only in computational peak-calling [1]; this bias leads to prediction of ‘new’ binding sites that result directly from this artifact, as demonstrated in the cross-validation results [2]. While the GraphProt prediction algorithm performs worse when terminating features are excluded from its feature set, it still benefits from having the ‘terminating G’ in the peaks and may also benefit from adjacent positions that are due to enzyme specificity. This finding implies that the observed bias cannot be easily removed computationally; distinguishing between the true protein binding preferences and the enzyme specificity may be impossible when the peaks are determined by both. Knowing the enzyme specificity in advance may allow us to deconvolve protein-binding signal from enzyme specificity and thereby accurately call unbiased peaks.

Thus, to solve the specific bias of ‘terminating G’s’ in CLIP data, we call for an appropriate experimental control. Previous reported controls accounted for RNA expression levels, but no control measured the cleaving preference of the enzyme [3]. The experiment should measure the enzyme specificity without the presence of an RNA-binding protein. Having such a control will allow assignment of prior cleaving probabilities to genomic-loci in the peak-calling (as done in Uren *et al*. [1] for other co-variants) and identification of unbiased binding sites. Such controls will lead to better algorithms to predict protein RNA-binding and thus more accurate prediction of new binding sites.

## Acknowledgments

This study was supported by NIH grant R01GM081871.

## Supplementary Methods

### Datasets

We downloaded 24 CLIP experiments from the GraphProt website (http://www.bioinf.uni-freiburg.de/Software/GraphProt/). The peaks were originally downloaded from the doRiNA database [4]. Control sequences were extracted from nearby regions on the same transcript (for details, see [2]).

For model training and evaluation, we used the sequences with three different flank length, denoted by *peaks±length*: *peaks+150*, *peaks* and *peaks-1*. In *peaks+150*, each sequence is flanked by 150bp on both sides for accurate structure prediction. In *peaks* flanks were removed to make the impact of the last nucleotide more visible. The removal of the last nucleotide showed how the ‘terminating G’ improved the accuracy of the models.

### Nucleotide distribution

For each experiment in the original GraphProt study we calculated the frequency of terminating G’s (the frequency of G’s at the end of a peak out of all peaks). In addition, we calculated the frequency of non-G nucleotides in the peaks and controls.

### Running GraphProt

For each dataset, we ran the algorithm as advised by the authors. For each experiment, 4 files were used: positives and controls for parameter optimization and positives and controls for training. We used the files ending with ‘ls.positives.fa’ and ‘ls.negatives.fa’ for parameter optimization and ‘train.positives.fa’ and ‘train.negatives.fa’ for model training.

We first optimized the parameters using the command line “perl GraphProt.pl – action ls –fasta <opt_peaks> -negfasta <opt_control>”. Following this procedure, we used the optimal values of epochs, lambda, R, D, bitsize and abstraction for 10-fold cross-validation. The command line: “perl GraphProt.pl –action cv – epochs <epochs> -lamba <lambda> -R <R> -D <D> -bistize <bistize> -fasta <non_opt_peaks> -negfasta <non_opt_control>”. We extracted the AUC value from the results. To train a model and get the feature weights, we trained a model on the ‘train.positive.fa’ and ‘train.negatives.fa’ files with the optimized parameters. To get the association between feature ID numbers and sequences we used the ‘-V’ verbose option in EDeN, run as part of GraphProt.pl script. RNAshapes version 2.1.6 was used to comply with its invocations in GraphProt scripts [5].

### P-value calculation

P-values in the manuscript were calculated using Wilcoxon rank-sum test. wilcox.test function in R was used with “paired=TRUE” and “alternative=two.sided”.

### GraphProt Modification

We modified GaphProt feature generation to exclude features encoding nucleotide k-mers at the end of the sequence. A feature is a pair of sub-graphs at distance d, each of radius r. If the size of one of the sub-graphs was smaller than r+1, we removed the feature from the feature set.

**Figure S1.**
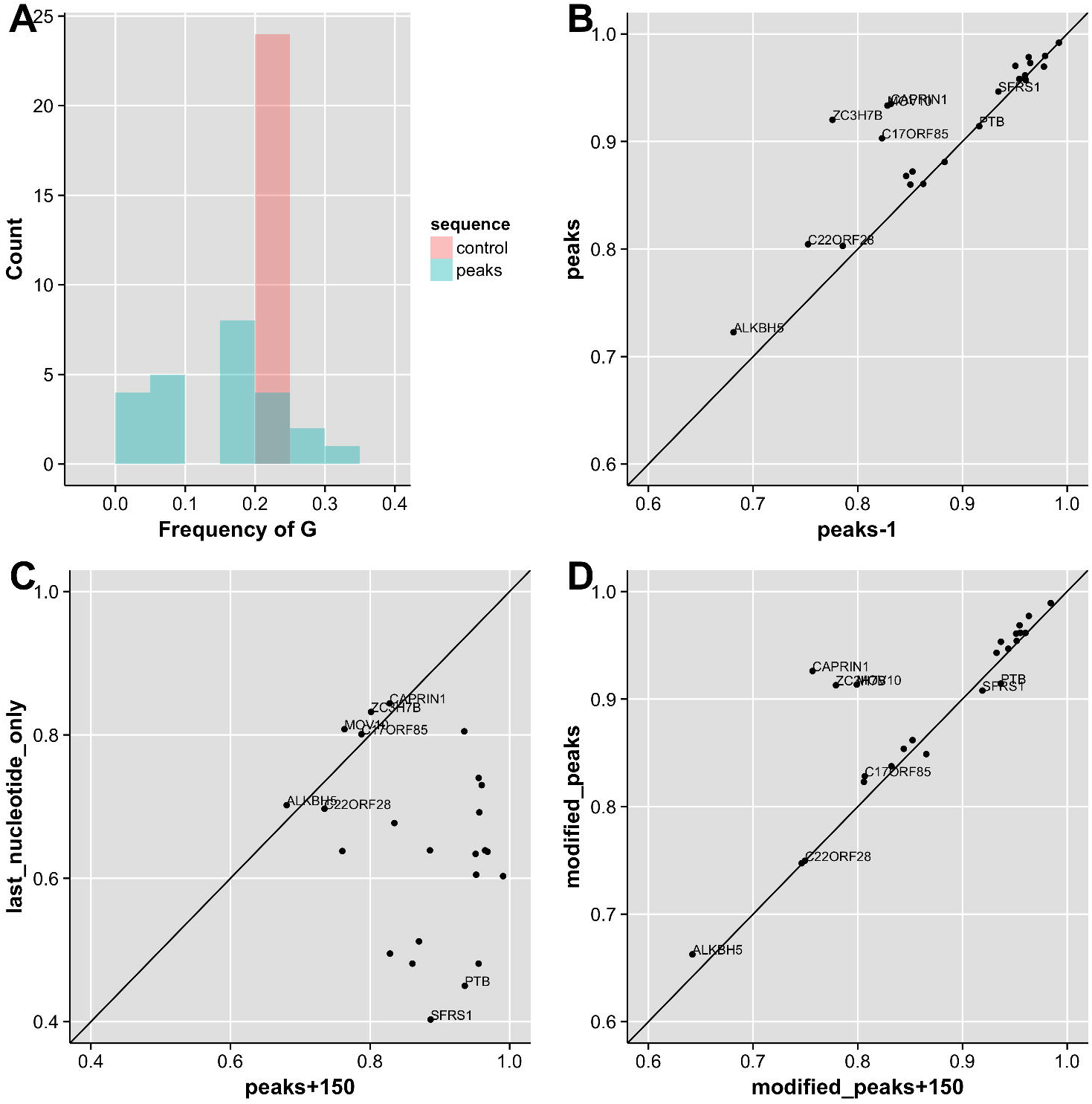
Additional sequence biases in CLIP experiments and their effect on binding models learned in downstream analysis. A) In most CLIP experiments the frequency of G’s is much lower in the peaks than in the control sequences. The histogram is over the frequency of G’s. B) GraphProt performs significantly better on CLIP experiments without the flanks but with the last nucleotide than without it due to the terminating G’s (see Table 1). C) For some experiments, the AUC achieved by ranking G-terminated sequences is even higher that achieved by GraphProt. In most experiment it is higher than random (0.5). D) Modified GraphProt performs significantly better on peaks without the flanks despite the exclusion of terminating nucleotides features.

## References

1. Uren PJ, Bahrami-Samani E, Burns SC, Qiao M, Karginov F V., Hodges E, Hannon GJ, Sanford JR, Penalva LOF, Smith AD: Site identification in high-throughput RNA-protein interaction data. Bioinformatics 2012, 28:3013–3020.

2. Maticzka D, Lange SJ, Costa F, Backofen R: GraphProt: modeling binding preferences of RNA-binding proteins. Genome Biol 2014, 15:R17.

3. Kishore S, Jaskiewicz L, Burger L, Hausser J, Khorshid M, Zavolan M: A quantitative analysis of CLIP methods for identifying binding sites of RNA-binding proteins. Nat Methods 2011, 8:559–64.

4. Anders G, Mackowiak SD, Jens M, Maaskola J, Kuntzagk A, Rajewsky N, Landthaler M, Dieterich C: doRiNA: a database of RNA interactions in posttranscriptional regulation. Nucleic Acids Res 2012, 40(Database issue):D180-6.

5. Steffen P, Voss B, Rehmsmeier M, Reeder J, Giegerich R: RNAshapes: an integrated RNA analysis package based on abstract shapes. Bioinformatics 2006, 22:500–3.

